# High-Yield Production of Cytotoxic Actinomycin D from Endophytic *Streptomyces parvulus* AL036 through Nutritional Optimization

**DOI:** 10.1101/2025.09.24.678420

**Authors:** Thongchai Taechowisan

## Abstract

Endophytic actinomycetes are bacteria that live inside plant tissues without causing harm to the plant, producing various bioactive compounds. This study aimed to isolate and characterize an endophytic actinomycete from *Alpinia galanga* roots, and to investigate its bioactive compounds. A *Streptomyces* strain designated AL036 was isolated and identified as *Streptomyces parvulus* through morphological, biochemical, and 16S rDNA sequence analysis. Similar to *Streptomyces parvulus* Tc022, this strain exhibited actinomycin D as its major bioactive compound. Media optimization experiments and cytotoxic activity against non-cancerous and cancer cell lines were carried out. The starch casein medium was identified as a medium supporting significantly higher actinomycin D production (103.67 mg/L) compared to the initial medium (ISP-2). Further optimization revealed that a specific carbon-to-nitrogen ratio (20 g/L soluble starch and 2 g/L casein) maximized production (199.33 mg/L). Sugar supplementation did not enhance production but influenced mycelial growth. The purified actinomycin D exhibited cytotoxicity against HeLa cells and LLC-MK2 cells, with IC_50_ values of 10.22 ± 2.53 µg/ml and 17.78 ± 3.97 µg/ml, respectively. This study isolated a new *Streptomyces parvulus* AL036, with efficient actinomycin D production similar to Tc022. Media optimization significantly improved production, highlighting the importance of carbon and nitrogen sources. Both crude extract and purified actinomycin D showed cytotoxicity and selectivity for some cancer cells.

## 1. INTRODUCTION

*Alpinia galanga*, a well-known medicinal herb, harbors beneficial endophytic actinomycetes— tiny bacteria that live within the plant in a fascinating, co-evolved partnership [1]. This symbiotic relationship enables the production of unique chemicals that bolster the plant’s growth, enhance its stress response, and even ward off disease, making herbaceous plants like galangal ideal hosts for these helpful microorganisms [2]. Scientists are actively exploring these endophytic actinomycetes as potential sources for new medicines and crop improvements, believing they may hold keys to a healthier future. Given the diverse bacterial population within galangal, there’s considerable interest in discovering further hidden gems.

Among endophytic actinomycetes, *Streptomyces* is the dominant genus [3]. A well-studied example is *Streptomyces parvulus*, renowned for producing actinomycin D, a powerful antibiotic. Actinomycins are a family of chromopeptide lactone antibiotics, with actinomycin D being the most extensively studied member. Produced by various *Streptomyces* species as part of a mixture, actinomycin D has demonstrated clinical efficacy in cancer treatment by inhibiting the cell cycle. This class of over 30 naturally occurring antibiotics shares a common phenoxazone chromophore, differing only in the amino acid composition of their peptide side chains [4]. While *Streptomyces* species are primary producers [5], *Micromonospora* has also been reported to synthesize actinomycins as part of a complex mixture [6-8]. The antimicrobial mechanism of action for actinomycin D involves binding to DNA and subsequently inhibiting RNA synthesis [9]. Historically significant as the first clinically utilized antibiotic in 1954, actinomycin D remains a valuable tool in both oncology and biochemical/molecular biological research [10].

In light of increasing antibiotic resistance, endophytic actinomycetes represent a promising reservoir of novel antibiotic compounds, potentially offering an alternative to current pharmaceutical production strategies. Our prior work, for instance, isolated *Streptomyces parvulus* Tc022 from *Alpinia galanga* root tissues, which was found to produce actinomycin D as a major compound [11], exhibiting a non-optimized yield of 66.31 mg/L when cultured in International *Streptomyces* Project medium 2 (ISP-2). While promising, it’s crucial to acknowledge the inherent challenges in transitioning from laboratory-scale production to industrial-scale manufacturing. Such a transition often involves optimizing not just the biological yield but also factors like bioreactor design, nutrient delivery, waste management, and purification processes to ensure cost-effectiveness and consistent quality at a much larger scale.

To further investigate the persistent endophytic association between actinomycetes and their plant hosts, this study aimed to re-isolate and characterize an endophytic actinomycete from the same *Alpinia galanga* roots in the same location again. A *Streptomyces* strain, designated AL036, was successfully isolated and identified as *Streptomyces parvulus* through morphological, biochemical, and 16S rDNA sequence analysis. Crucially, similar to *Streptomyces parvulus* Tc022, this re-isolated strain exhibited actinomycin D as its major bioactive compound, providing unique evidence of remarkable genetic and metabolic consistency of this endophytic strain over a seventeen-year period. Furthermore, this study explored the optimization of culture media to significantly enhance actinomycin D production by strain AL036. A novel Starch Casein (SC) medium was specifically developed and optimized, resulting in a substantial improvement in actinomycin D yield. Interestingly, supplementation with monosaccharides (glucose, galactose, fructose) was found to repress actinomycin D biosynthesis while promoting mycelial growth, likely due to carbon catabolite repression. The anticancer activity of the purified actinomycin D was confirmed against HeLa cells, demonstrating its continued biological efficacy. These findings aim to improve the foundational understanding necessary for future industrial applications and validate the potential of consistently re-isolated endophytic strains.

## 2. MATERIALS AND METHODS

### 2.1. Sample preparation

*Alpinia galanga* roots were collected from a site located at Baan Suan Takhrai, Nakhon Pathom, Thailand (13.82859°N, 100.04352°E). To isolate actinomycetes, the samples were cut into 5 x 5 mm^2^ pieces, thoroughly washed, and then subjected to a multi-step sterilization process. This process involved rinsing with Tween 20 solution, sodium hypochlorite, and ethanol to remove surface contaminants. Finally, the sterilized tissue pieces were dried aseptically in a laminar flow cabinet (Esco Scientific, PA, USA). The surface-sterilized plant tissues were plated onto humic acid-vitamins (HV) agar [12]. To prevent fungal and yeast growth, 100 µg/ml of cycloheximide and nystatin were added to the agar. These plates were incubated at 32°C for 3 weeks. Colonies exhibiting characteristic actinomycete morphologies were picked and transferred to fresh plates containing International *Streptomyces* Project medium 2 (ISP-2). Among the total of 35 actinomycete isolates evaluated for their ability to inhibit bacterial growth, the isolate designated AL036 showed potent activity against the tested bacteria. This isolate was selected for identification and further analysis.

### 2.2. Identification of AL036 strain

Morphological properties were observed with a scanning electron microscope (TESCAN Mira3, Czech Republic). All chemicals and media components were purchased from Hi-media Laboratories (Mumbai, India) and used to investigate the physiological characteristics. Chemotaxonomy and 16S rRNA gene sequence analysis were carried out using the method described by Taechowisan et al. [13].

### 2.3. Growth, production media, and metabolite extraction

Starter cultures in 250-ml Erlenmeyer flasks containing 25 ml of growth medium were inoculated with a 1 ml suspension of AL036 strain (7.0 × 10^7^ spores/ml). These flasks were incubated at 32°C, shaken at 200 rpm for 48 hours. After this period, the starter cultures were transferred to 3 L Erlenmeyer flasks containing 1.5 L of five different media and incubated at 32°C, shaken at 200 rpm for 21 days for actinomycin D production. The five media used were presented (Table 1).

**Table 1.** Medium components used for Actinomycin D production.

| Medium | Components (g/L) |
| --- | --- |
| Starch Casein (SC) | Soluble starch (10), casein (1), potassium nitrate (2), sodium chloride (2), dipotassium hydrogen phosphate (2), magnesium sulfate heptahydrate (0.05), calcium carbonate (0.02), ferrous sulfate heptahydrate (0.01) |
| Bennett's | Glucose (10), N-Z amine type A (2), yeast extract (1), beef extract (1) |
| SM3 | Glucose (10), peptone (5), tryptone (3) |
| Mueller-Hinton | Acid hydrolysate of casein (17.5), beef extract (2), soluble starch (1.5) |
| International<br>*Streptomyces*<br>Project 2 (ISP-2) | Malt extract (10), yeast extract (4), glucose (4) |

The final volume of all culture media was adjusted to 1000 ml with distilled water, and the pH was adjusted to between 7.0 and 7.2. Following incubation, the cultures were filtered twice through cotton filters. Ethyl acetate extraction of the supernatants followed by vacuum concentration yielded a crude metabolite extract.

### 2.4. Purification and chemical characterization

The crude metabolite was subjected to column chromatography using silica gel 60 (Merck, 0.040-0.063 mm) and CH_2_Cl_2_/MeOH/H_2_O (20:3:1) as the eluent to yield actinomycin D as a red-orange solid. The production of actinomycin D was quantified across various culture media. Thin-layer chromatography (TLC) was used to assess the composition and purity of the mixtures by separating their constituent components. The purified actinomycin D was characterized by determining its melting point, electrospray ionization high-resolution mass spectrometry (ESI-HRMS), and ^1^H- and ^13^C-nuclear magnetic resonance (NMR) spectral data.

### 2.5. Optimization of actinomycin D production

SC medium was identified as supporting the highest actinomycin D production and was designated as the basal medium for further optimization. A two-factor experiment utilizing level factors was designed to optimize the concentrations of soluble starch and casein for enhanced actinomycin D yield. Each factor, soluble starch and casein, was investigated at three different levels: 5, 10, and 20 g/L for starch and 0.5, 1.0, and 2.0 g/L for casein. These levels were combined in a full factorial design. The cultivation conditions and procedures for crude metabolite extraction were identical to those previously described. To further enhance actinomycin D production, the medium optimized for its highest yield was supplemented with glucose, galactose, and fructose at concentrations of 0.5% and 1% (w/v) to assess their impact on actinomycin D titer and dry weight.

### 2.6. Determination of the cytotoxicity activity of the crude extract and purified compound

The *in vitro* cytotoxicity of the crude extract and the purified actinomycin D was determined using the MTT assay [14]. HeLa cells, a human cervical carcinoma cell line, were exposed to a concentration range of 1-512 µg/ml of both the extract and the compound. To evaluate the compound’s selectivity towards cancer cells, a non-cancerous LLC-MK2 cell line was included in the assay. IC_50_ values, representing the concentration inhibiting 50% of cell growth, were calculated using non-linear regression analysis of the dose-response curves. Selectivity indices (SI) were calculated as the ratio of the IC_50_ values obtained in the LLC-MK2 and HeLa cells. Higher SI values indicate greater selectivity of the compound for cancer cells, with minimal cytotoxicity towards healthy cells. Commercial actinomycin D and doxorubicin hydrochloride (Thermo Fisher Scientific, Massachusetts, USA) served as positive controls for cytotoxicity.

### 2.7. Statistical analyses

All experiments were performed in triplicate. Data were expressed as means ± standard deviations (SD). Statistical analyses for comparing treatment effects (e.g., actinomycin D production in different media or optimized conditions) were performed using one-way Analysis of Variance (ANOVA), followed by Duncan’s multiple range tests for post-hoc comparisons. All statistical computations were carried out using SPSS for Windows version 11.01 (SPSS Inc., Chicago, IL, USA). A *p*-value of α 0.05 was considered to indicate statistical significance.

## 3. RESULTS

A total of 35 strains were isolated from *Alpinia galanga* roots. Based on screening for antibacterial activity using the soft-agar overlay method, one actinomycete isolate (AL036) showed potent activity against the tested bacteria. This isolate was selected for identification and further analysis. The colony of AL036 was yellow-brown with orange-red soluble pigment. Scanning electron microscopy (SEM) revealed the formation of rectiflexibile chains containing smooth-surfaced spores on lateral branches of aerial hyphae (Figure 1). LL-diaminopimelic acid was detected in the cell extract. The physiological and biochemical properties are presented (Table 2). Based on these characteristics, the strain was identified as a member of the genus *Streptomyces*. The strain was further identified by 16S rRNA gene sequence analysis (GenBank Accession No. PP972269). NCBI BLAST search analysis with the 16S rRNA gene showed that the sequence was 99.68% and 100% similar to the sequence of *Streptomyces parvulus* SDMRI3 and *Streptomyces parvulus* Tc022, respectively. The phylogenetic position of *Streptomyces parvulus* AL036 is presented (Figure 2).

**Table 2.**
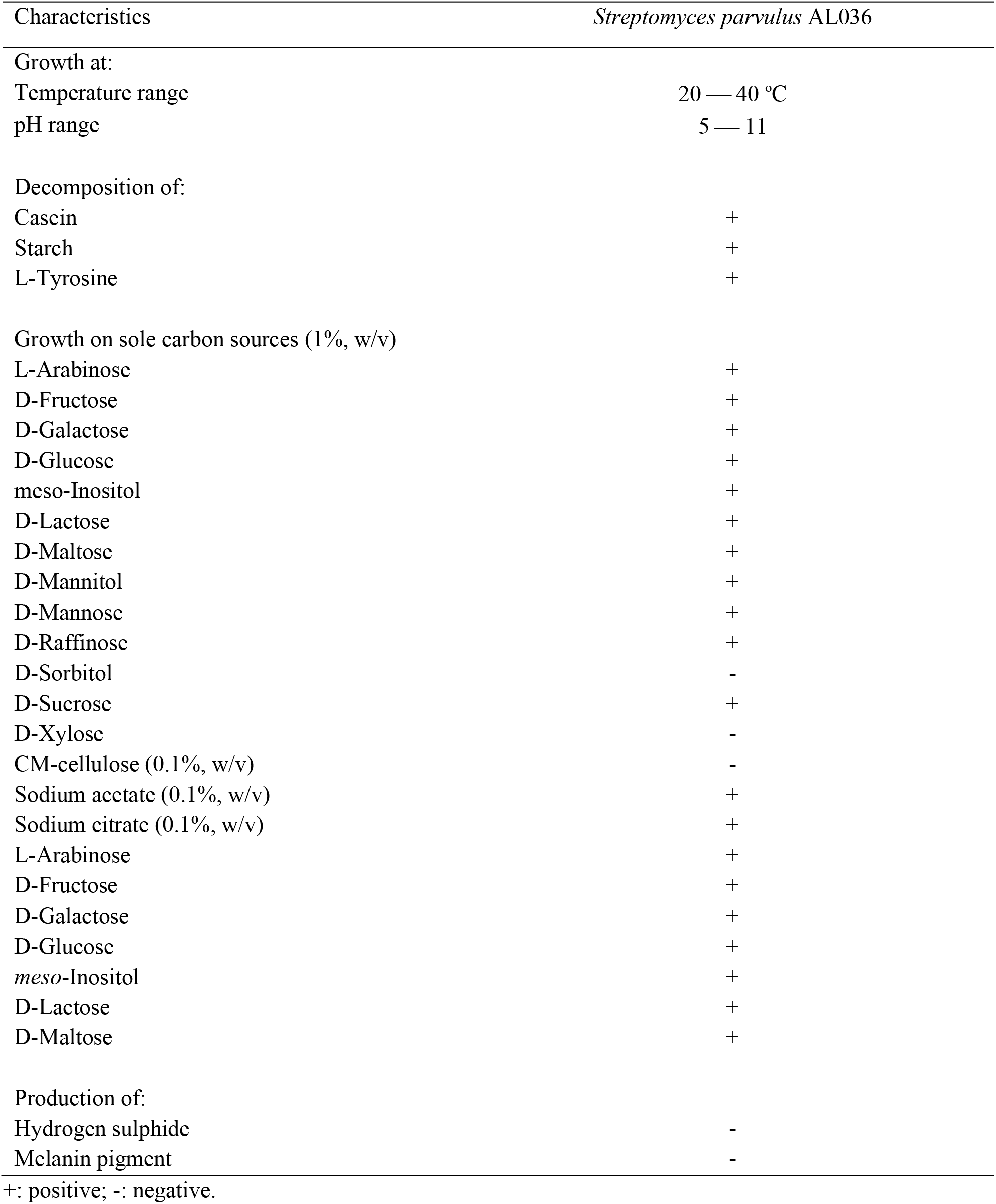
Characteristics of strain *Streptomyces parvulus* AL036.

| Characteristics | *Streptomyces parvulus* AL036 |
| --- | --- |
| Growth at: |  |
| Temperature range | 20 — 40 °C |
| pH range | 5 — 11 |
| Decomposition of: |  |
| Casein | + |
| Starch | + |
| L-Tyrosine | + |
| Growth on sole carbon sources (1%, w/v) |  |
| L-Arabinose | + |
| D-Fructose | + |
| D-Galactose | + |
| D-Glucose | + |
| meso-Inositol | + |
| D-Lactose | + |
| D-Maltose | + |
| D-Mannitol | + |
| D-Mannose | + |
| D-Raffinose | + |
| D-Sorbitol | - |
| D-Sucrose | + |
| D-Xylose | - |
| CM-cellulose (0.1%, w/v) | - |
| Sodium acetate (0.1%, w/v) | + |
| Sodium citrate (0.1%, w/v) | + |
| L-Arabinose | + |
| D-Fructose | + |
| D-Galactose | + |
| D-Glucose | + |
| meso-Inositol | + |
| D-Lactose | + |
| D-Maltose | + |
| Production of: |  |
| Hydrogen sulphide | - |
| Melanin pigment | - |

+: positive; -: negative.

**Figure 1.**
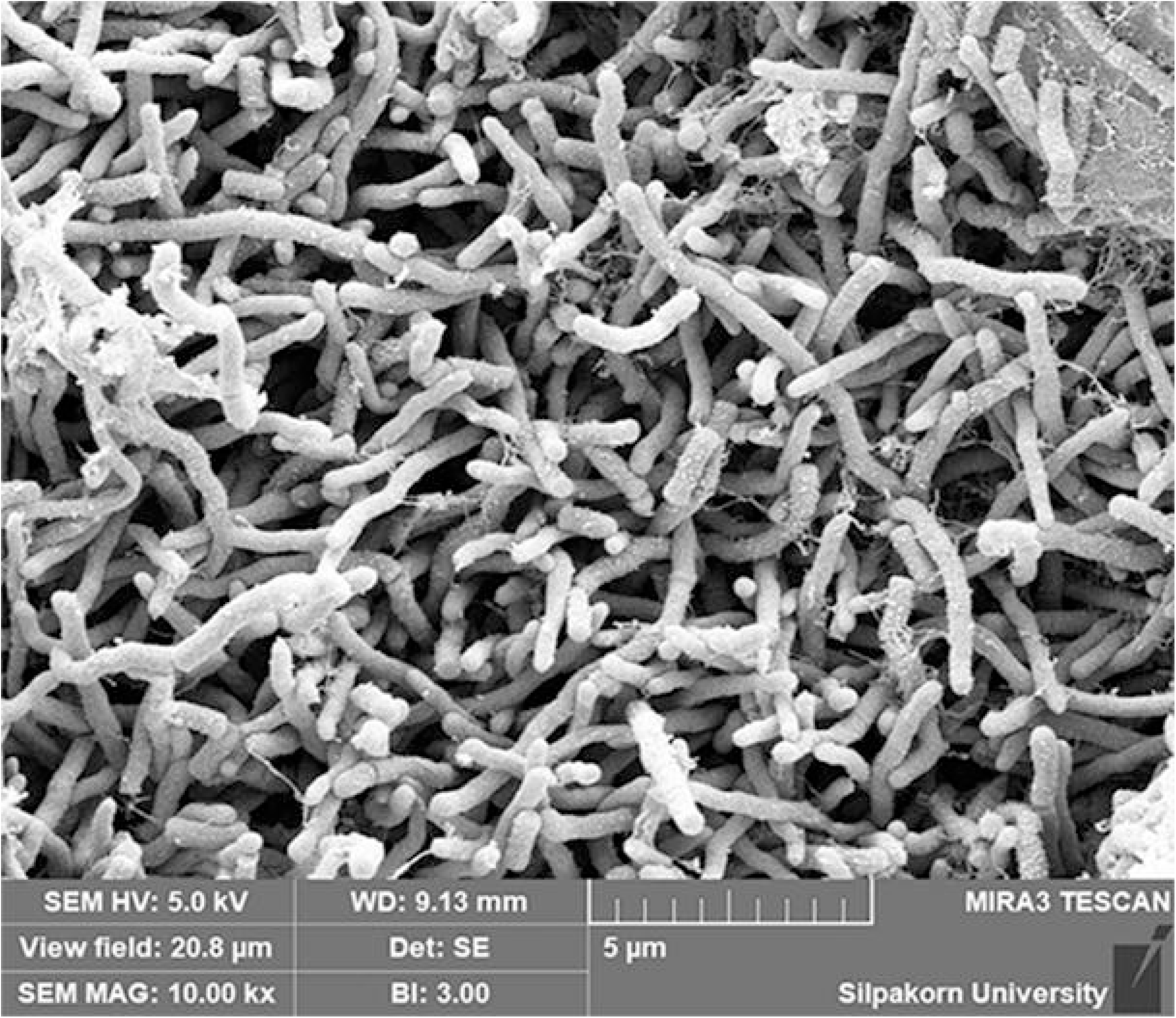
Scanning electron micrograph of *Streptomyces parvulus* AL036 after 21 days cultivation on ISP-2 medium at 32°C.

**Figure 2.**
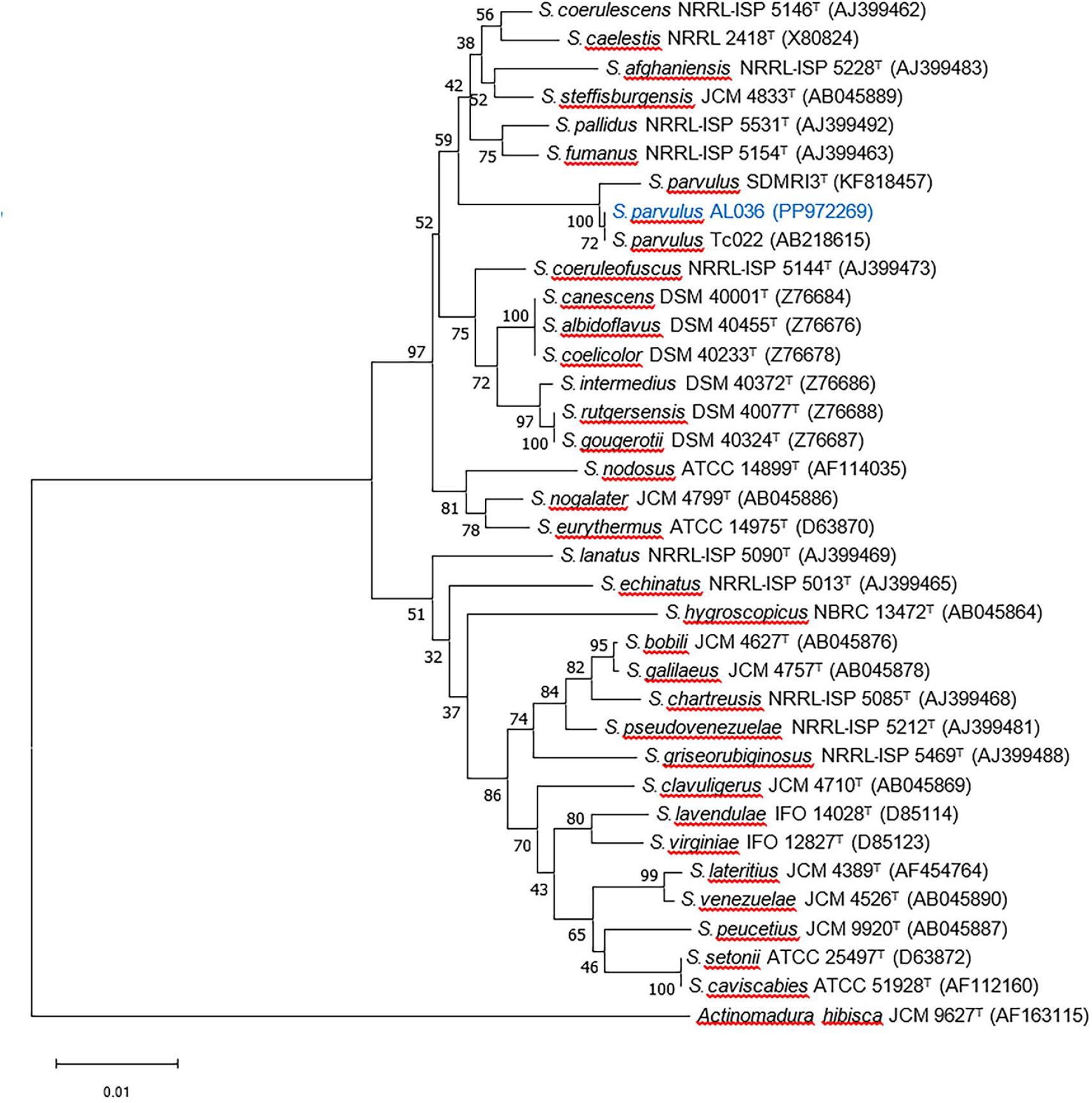
Neighbor-joining tree base on partial 16S rRNA gene sequences showing relationships among *Streptomyces parvulus* AL036 (PP972269) and related menbers of the genus *Streptomyces*. *Actinomadura hibisca* JCM 9627^T^ was used as outgroup. Bootstrap values greater than 50% are highlighted at the nodes (1000 replications).

After isolation, the major compound was available in sufficient amounts for further studies. The ^1^H-NMR, ^13^C-NMR, and mass spectral data were identical to those of actinomycin D (Supplementary Materials: Figures S1 – S3). In the experiment conducted for isolation of actinomycin D in different media, it was found that *Streptomyces parvulus* AL036 could produce actinomycin D at yields ranging from 56.67 to 103.67 mg/L across various culture media. In SC medium, the growth of *Streptomyces parvulus* AL036 and the production of actinomycin D showed significant advantages (Table 3). Specifically, the production of actinomycin D in SC medium reached 103.67 ± 2.52 mg/L at 32°C for 21 days, which was statistically higher than the actinomycin D yield of 56.67 ± 4.51 mg/L obtained in ISP-2 medium (*p* α 0.05, based on one-way ANOVA and Duncan’s multiple range test).

**Table 3.** Biomass and actinomycin D production in different culture media.

| Culture media | Actinomycin D (mg/L)* |
| --- | --- |
| Starch Casein medium | 103.67 ± 2.52<sup>a**</sup> |
| Bennett's medium | 80.67 ± 4.51<sup>b</sup> |
| SM3 medium | 71.33 ± 4.73<sup>c</sup> |
| Mueller-Hinton medium | 84.33 ± 5.03<sup>b</sup> |
| ISP-2 medium | 56.67 ± 4.51<sup>d</sup> |

\*Cultivation at 32°C, shaken at 200 rpm for 21 days.
\*\*Different letters indicated statistically significant differences ( $p < 0.05$ ).

The SC medium was selected to evaluate the quantitative influence of carbon and nitrogen (C&N) sources upon actinomycin D production by *Streptomyces parvulus* AL036. The concentrations of soluble starch and casein were varied in recombination to provide different C&N concentrations in the medium. The results of these experiments are shown (Table 4). Higher production of actinomycin D (199.33 mg/L) was achieved at C&N concentrations of 20 g/L for soluble starch and 2 g/L of casein, which was statistically significant compared to other concentrations (*p* α 0.05, based on one-way ANOVA and Duncan’s multiple range test). This optimized C&N starch-casein medium was then used to investigate the effect of various sugar supplementations on actinomycin D production. Supplementation of sugars (glucose, galactose, and fructose) to the complete C&N starch-casein medium did not significantly enhance actinomycin D production compared to the unsupplemented medium (Table 5). However, the presence of these sugar supplements did influence mycelial dry weight.

**Table 4.** Effect of several C&N concentrations on actinomycin D production by *Streptomyces parvulus* AL036 grown in a starch -casein medium.

| Starch (g/L) | Casein (g/L) | C/N ratios | Actinomycin D (mg/L)* |
| --- | --- | --- | --- |
| 5 | 0.5 | 10 | 78.33 ± 5.03<sup>abcde**</sup> |
|  | 1 | 5 | 76.67 ± 3.05<sup>abc</sup> |
|  | 2 | 2.5 | 79.67 ± 2.52<sup>abcde</sup> |
| 10 | 0.5 | 20 | 88.33 ± 2.52<sup>acde</sup> |
|  | 1 | 10 | 89.33 ± 3.21<sup>acde</sup> |
|  | 2 | 5 | 120.67 ± 11.50<sup>f</sup> |
| 20 | 0.5 | 40 | 171.67 ± 7.09<sup>g</sup> |
|  | 1 | 20 | 178.33 ± 4.04<sup>g</sup> |
|  | 2 | 10 | 199.33 ± 11.67<sup>h</sup> |

\*Cultivation at 32°C, shaken at 200 rpm for 21 days.
\*\*Different letters indicated statistically significant differences ( $p < 0.05$ ).

**Table 5.** Effect of sugar supplement in complete C&N starch -casein medium on mycelial dry weight and actinomycin D production by *Streptomyces parvulus* AL036.

| Sugar supplement | Mycelial dry weight (g/L)* | Actinomycin D (mg/L)* |
| --- | --- | --- |
| 0.5% Glucose | 30.35 ± 1.05<sup>a**</sup> | 106.67 ± 6.03<sup>acd**</sup> |
| 1.0% Glucose | 32.59 ± 0.68<sup>a</sup> | 128.67 ± 15.31<sup>bf</sup> |
| 0.5% Galactose | 29.48 ± 1.10<sup>a</sup> | 95.33 ± 10.21<sup>acde</sup> |
| 1.0% Galactose | 31.50 ± 0.90<sup>a</sup> | 109.33 ± 7.57<sup>cf</sup> |
| 0.5% Fructose | 29.60 ± 1.13<sup>a</sup> | 84.33 ± 8.02<sup>ce</sup> |
| 1.0% Fructose | 31.41 ± 0.88<sup>a</sup> | 126.67 ± 12.66<sup>bdf</sup> |
| Non sugar supplement | 24.59 ± 1.84<sup>b</sup> | 199.33 ± 11.68<sup>g</sup> |

\*Cultivation at 32°C, shaken at 200 rpm for 21 days.
\*\*Different letters indicated statistically significant differences ( $p < 0.05$ ).

The crude extract and purified actinomycin D were assessed for cytotoxicity using a non-cancerous rhesus kidney monkey cell line (LLC-MK2) and a human cervical carcinoma cell line (HeLa). They exhibited high cytotoxicity, with IC_50_ values ranging from 10.22 to 19.52 µg/ml for HeLa cells. The crude extract and purified actinomycin D showed moderate cytotoxicity against the non-cancerous cell line (LLC-MK2), with IC_50_ values of 32.21 and 17.78 µg/ml, respectively. The selectivity indices (SI) of crude extract, purified actinomycin D, and commercial actinomycin D against the cancer cells were significantly lower compared to doxorubicin hydrochloride (*p* α 0.05, based on one-way ANOVA and Duncan’s multiple range test) (Table 6).

**Table 6.**
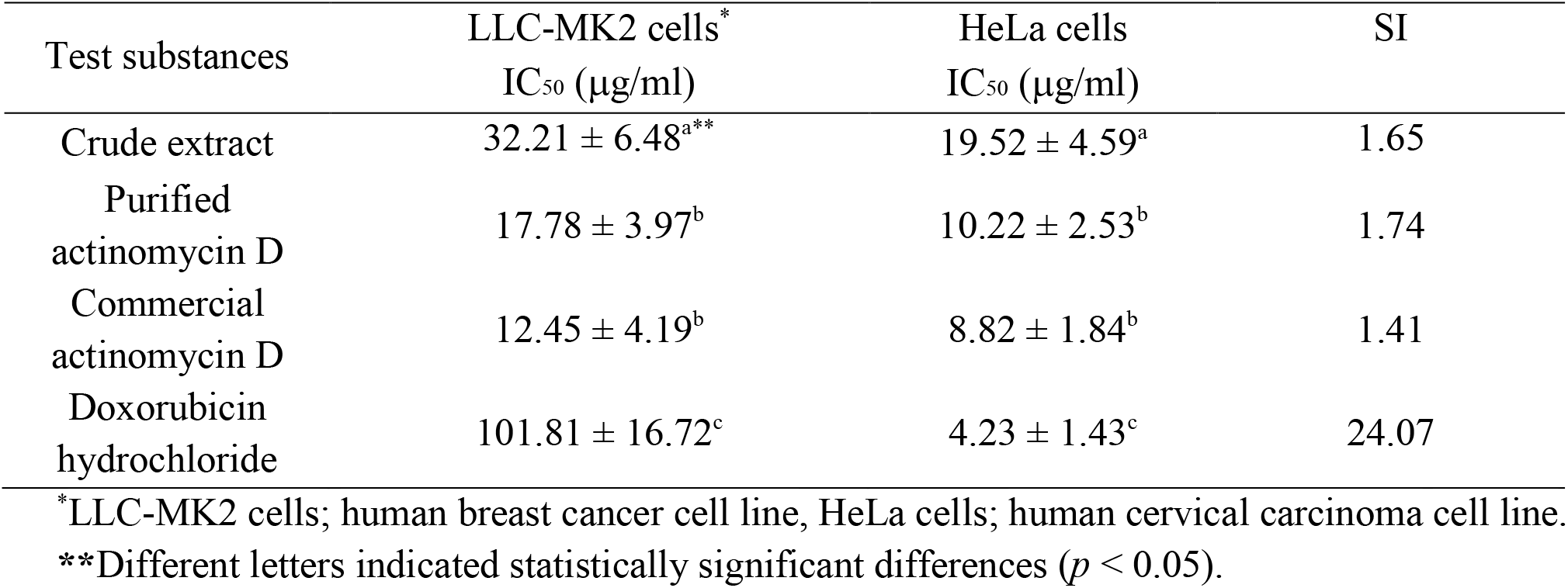
IC_50_ values and selectivity indices (SI) of crude extract and purified actinomycin D against non-cancerous and cancer cell lines.

## 4. DISCUSSION

*Streptomyces parvulus* AL036, isolated from *Alpinia galanga* root tissues, showed strong similarities in morphological, physiological, biochemical characteristics, and 16S rDNA sequence to *Streptomyces parvulus* Tc022. This consistency suggests that AL036 and Tc022 might be the same strain, indicating its persistent presence within the *Alpinia galanga* root niche over seventeen years since its initial isolation in 2006 [11]. Our study successfully identified and characterized actinomycin D, a known anticancer antibiotic, produced by *S. parvulus* AL036 from *Alpinia galanga*. The compound’s identity and bioactivity were confirmed through ESI-HRMS and NMR data.

Actinomycin D yields can vary substantially among different *Streptomyces* species. For instance, *S. parvulus, S. felleus*, and *S. regensis* have been reported to produce 152, 20, and 12 mg/ml of actinomycin D, respectively, with *S. parvulus* known for almost exclusive actinomycin D production [15, 16]. However, most strains typically produce only small quantities. Our study focused on optimizing the production of actinomycin D by *S. parvulus* AL036, starting with the identification of SC medium as the most favorable initial condition. Optimizing carbon and nitrogen (C&N) nutrient concentrations and monosaccharide supplements is crucial for maximizing actinomycin D yield in both laboratory and industrial fermentation processes, as these parameters significantly impact cell growth and production stability.

This study’s findings on actinomycin D production by *Streptomyces parvulus* AL036 present some interesting comparisons with prior research, particularly the work of Sousa et al. [16]. While Sousa et al. utilized a different strain (*S. parvulus* DAUFPE 3124) and a distinct medium (soy milk-based with glucose and CaCO_3_), our research employed strain AL036 and a novel SC medium based on soluble starch and casein. The optimization strategies also differed; Sousa et al. primarily investigated substituting glucose with other sugars, especially fructose, whereas we systematically optimized the base SC medium by manipulating C&N concentrations before examining the impact of sugar supplementation.

These differences in approach led to contrasting results regarding the effect of sugars. Sousa et al. observed increased actinomycin D production with fructose compared to glucose, while our study found that supplementing the optimized SC medium with glucose, galactose, or fructose actually inhibited actinomycin D production, despite stimulating mycelial growth. This suggests that our optimized SC medium already provided an optimal carbon source balance, and additional sugars disrupted this balance.

When comparing actinomycin D yields, Sousa et al. reported substantially higher yields (530-635 mg/L) than the 199.33 mg/L achieved with our optimized SC medium. This discrepancy in yield is a critical point that warrants further discussion. Several factors could contribute to the lower yields observed in our study: 1) Strain Variation: Even within the same species, different *Streptomyces parvulus* strains can possess distinct genetic backgrounds and metabolic capabilities, leading to varying levels of secondary metabolite production. The DAUFPE 3124 strain used by Sousa et al. might inherently be a stronger producer than our AL036 strain under their specific fermentation conditions. 2) Media Composition and Optimization: While we optimized our SC medium, the fundamental composition might still differ significantly from the soy milk-based medium of Sousa et al. The specific blend of nutrients, trace elements, and buffers in their medium could be more conducive to higher actinomycin D biosynthesis for their strain. Our finding that additional sugars inhibited production in our optimized SC medium suggests a delicate balance; their medium, or strain, might respond differently to sugar supplementation. 3) Fermentation Conditions: Beyond media composition, other fermentation parameters like initial pH, aeration rates, agitation speeds, and total fermentation time can profoundly impact product yield. Although both studies likely maintained optimal temperatures, subtle differences in these process parameters (e.g., specific bioreactor configurations or oxygen transfer rates) could account for significant variations in final productivity. For instance, *Streptomyces* species are obligate aerobes requiring optimal aeration for growth and metabolism [20]. While our study used a constant 32°C and 200 rpm shaking, previous work by Praveen and coauthors demonstrated a 356% increase in actinomycin D yield by optimizing both nutrient composition and bioreactor operation parameters (agitation and aeration) compared to shake flask cultures using a non-optimized medium [21]. This highlights that our current shake flask conditions, while optimized for media, may still limit the full production potential compared to advanced bioreactor systems.

In essence, while both studies explored actinomycin D production by *Streptomyces parvulus*, their use of different strains, media, and optimization strategies led to contrasting outcomes, especially concerning the effect of sugar supplementation. This underscores the importance of strain-specific optimization for maximizing secondary metabolite production and highlights the complex challenges involved in scaling up actinomycin D production from laboratory to industrial scales. Achieving commercially viable production levels for actinomycin D will likely necessitate further bioprocess optimization, potentially involving more sophisticated bioreactor designs and precise control over environmental parameters.

The observed inhibition of actinomycin D biosynthesis by glucose, galactose, or fructose supplementation in *Streptomyces parvulus* AL036 likely stems from carbon catabolite repression (CCR), a well-documented regulatory mechanism in microorganisms [17, 18]. This phenomenon involves the preferential utilization of readily metabolizable sugars (like glucose) over other carbon sources, often repressing the expression and/or activity of key enzymes involved in alternative metabolic pathways or secondary metabolite biosynthesis. Specifically, this study aligns with findings by Gallo and Katz [17], who observed significant (∼94%) glucose-mediated repression of phenoxazinone synthase. This enzyme catalyzes the formation of the phenoxazinone ring, a crucial building block of the actinomycin D molecule [19]. Thus, the presence of these supplementary sugars likely leads to reduced phenoxazinone synthase activity, thereby limiting actinomycin D synthesis.

Beyond the media optimization and yield comparisons, another significant aspect of this study is the re-isolation of *Streptomyces parvulus* AL036 from *Alpinia galanga* roots. This re-isolation, from the same geographical location as the original Tc022 strain after a seventeen-year interval, raises an intriguing question: does frequent or “fresh” isolation of endophytic *Streptomyces* strains lead to a greater production of secondary metabolites compared to long-term laboratory cultures? While our current study focused on optimizing the culture medium for the re-isolated strain, the persistent presence and consistent actinomycin D production capabilities of AL036, mirroring those of Tc022, suggest a stable endophytic relationship. However, it’s a widely discussed topic in microbial natural product discovery that freshly isolated strains often exhibit higher metabolic activity and a broader range of secondary metabolite production than strains maintained under laboratory conditions for extended periods. This phenomenon can be attributed to several factors: 1) Loss of Biosynthetic Potential: Over numerous subcultures in artificial laboratory media, microorganisms can experience genetic drift or epigenetic changes. Genes responsible for the biosynthesis of complex secondary metabolites, which might not be essential for survival in nutrient-rich lab environments, can be downregulated, mutated, or even lost. This leads to a decrease in the production of these valuable compounds. 2) Environmental Cues: In their natural endophytic habitat within the plant, *Streptomyces* strains are exposed to a complex interplay of signals from the host plant and other co-existing microorganisms. These environmental cues can act as triggers for the activation of silent or cryptic biosynthetic gene clusters, leading to the production of novel or higher quantities of known compounds. Laboratory conditions, even optimized ones, rarely fully replicate this intricate natural environment. 3) Selective Pressure: In nature, the production of secondary metabolites often provides a selective advantage, such as antimicrobial activity against competitors or protective effects for the host plant. This continuous selective pressure helps maintain the biosynthetic machinery. In a lab setting without such pressures, the energetic burden of producing these compounds might lead to their downregulation. Therefore, the re-isolation of AL036, while demonstrating consistent production, also opens avenues for future research. It would be highly valuable to conduct comparative studies directly assessing secondary metabolite profiles and yields (e.g., through metabolomics approaches) between newly isolated endophytic *Streptomyces* strains and their long-term cultured counterparts. Such studies could provide crucial insights into strategies for maintaining and even enhancing the biosynthetic potential of these valuable microorganisms for pharmaceutical and industrial applications.

## 5. CONCLUSION

This study successfully re-isolated a *Streptomyces parvulus* strain, designated AL036, from *Alpinia galanga* roots collected from the same location as a previous study conducted in 2006. AL036 displayed morphological, physiological, and biochemical characteristics, a 16S rDNA sequence, and actinomycin D production capabilities remarkably similar to those of *Streptomyces parvulus* Tc022. These findings suggest the possibility that these endophytic actinomycetes might represent identical strains persistently residing within their host root (*Alpinia galanga*) niche over a seventeen-year period. Notably, *Streptomyces parvulus* AL036 produced actinomycin D, a bioactive metabolite exhibiting anticancer activity against HeLa cells. SC medium was identified as the most favorable for actinomycin D production. Further optimization of the SC medium identified a combination of 20 g/L soluble starch and 2 g/L casein as the most effective, resulting in a significant improvement in actinomycin D yield (from 103.67 mg/L to 199.33 mg/L). Interestingly, supplementation with 0.5 – 1.0% monosaccharides exhibited a repressive effect on actinomycin D production, likely due to carbon catabolite repression.

## 6. ACKNOWLEDGEMENTS

This study was financially supported by the seed grant SRIF-JRG-2568-08 from the Faculty of Science, Silpakorn University, Nakhon Pathom, Thailand.

## 7. PREPRINT SERVER NAME

A preprint version of this manuscript has been published on bioRxiv (MS ID#: BIORXIV/2025/644871). The authors undertake that they will update the preprint with a link to the final published version once accepted.

## 8. AUTHOR CONTRIBUTIONS

TT conceived and designed the experiments, performed the experiments, analyzed the data, prepared figures and tables, and prepared drafts of the manuscript. PK and TC conceived and designed the experiments, authored drafts of the manuscript, and evaluated bioactivity assays. WSP conceived and designed the experiments, purification, and structure elucidation of bioactive compounds.

## 9. CONFLICTS OF INTEREST

The authors report no financial or any other conflicts of interest in this work.

## 10. ETHICAL APPROVALS

This study does not involve human or animal subjects, and therefore does not require ethical approval.

## 11. USE OF ARTIFICIAL INTELLIGENCE (AI)-ASSISTED TECHNOLOGY

The authors declare that they have not used artificial intelligence (AI) tools for writing and editing the manuscript, and no images were manipulated using AI.

## Notes

### Competing Interest Statement

The authors have declared no competing interest.

